# Methylation-To-Expression Feature Models of Breast Cancer Accurately Predict Overall Survival, Distant-Recurrence Free Survival, And Pathologic Complete Response in Multiple Cohorts

**DOI:** 10.1101/187526

**Authors:** Jeffrey A. Thompson, Brock C. Christensen, Carmen J. Marsit

## Abstract

**Background:** Approaches that capitalize on the benefits of multi-omic data integration in invasive breast carcinoma to define prognostic biomarkers for precision medicine have been slow to emerge. In this work, we examined the efficacy of our methylation-to-expression feature model (M2EFM) approach to combining molecular and clinical predictors as part of a single analysis to create prognostic risk scores for overall survival, distant metastasis, and chemosensitivity.

**Methods:** Gene expression and DNA methylation values as well as clinical variables were integrated via M2EFM to build prognostic models of overall survival using 1028 breast tumor samples and further applied to external validation cohorts of 61 and 327 samples. Data-integrated prognostic models of distant recurrence-free survival and pathologic complete response were built using 306 samples and validated on 182 samples of external validation data. Additionally, we compared the discrimination and calibration of M2EFM models to other approaches.

**Results:** Despite different populations and assays, M2EFM models validated with good accuracy (C-index or AUC ≥ .7) for all outcomes in all validation data. M2EFM models had the most consistent performance overall and superior calibration, suggesting a greater likelihood of clinical utility. Finally, we demonstrated that M2EFM identifies functionally relevant genes, which could be useful in translating an M2EFM biomarker to the clinic.

**Conclusion:** M2EFM uses multiple levels of genomic data to infer disrupted regulatory patterns, thus providing a gene signature that connects loss of regulatory control with cancer prognosis.

**Funding:** The analyses described in this report were supported by NIH grants R01ES022222, P30CA138292, P30ES019776, and R01DE022772.

**Conflicts of Interest:** The authors declare no potential conflicts of interest.

## Background

The extreme heterogeneity of breast cancer makes one-size-fits all treatments difficult to achieve and although a number of targeted therapies are available (1) personalized therapies work only for a limited number of patients. Better methods are needed to rapidly identify patients than can benefit from specific treatments, reveal new targets for historically difficult to treat patients, and providing accurate prognoses to patients to enable better life planning.

Large scale collection of cancer genomic data, such as The Cancer Genome Atlas (TCGA) (2) and the International Cancer Genome Consortium (3), and expression repositories such as NCBI’s Gene Expression Omnibus (4) have enabled developments in prognostic and predictive cancer models. In addition to improved understanding of how cancer disrupts normal cellular function, such data have led to the development of biomarker panels, such as the PAM50 risk of recurrence score (rorS) (5) and the 21 gene Oncotype DX (6), which are both now used for clinical decision making in the treatment of breast cancer (7). Furthermore, to the extent that such tests are able to predict the occurrence of distant metastases, they may enable early prevention strategies to be put in place.

Despite such promising developments, the goals of precision medicine remain elusive due to specific limitations in molecular biomarker studies. These include limited understanding of the optimal genomic features to utilize, lack of validation on external data, and inconsistent use of best practices for creating biomarkers with the best chances of entering clinical practice (8, 9).

There has been growing interest in the integration of molecular and traditional clinicopathological markers to achieve higher levels of prognostic power and utility and small but significant gains in prognostic accuracy have been observed through this approach (10, 5). To the extent that these sources are independently prognostic, they should result in higher accuracy than either source alone and better understanding of prognostic features by revealing holistic relationships in the data. As yet, few prognostic or predictive models are designed to take advantage of multiple data types. In this work, we apply our own previously developed data integration approach for combining gene expression and DNA-methylation data to model the impact of regulatory disruption on breast cancer prognosis and prediction while also incorporating prognostic clinical variables (10). This study overcomes prior limitations by creating a model that can identify biologically plausible targets, using best practices in assessing the model (8, 9), and by validating the actual model (not just the gene signature) in multiple, external datasets, representing different demographics and disease distributions.

## Methods

### M2EFM

Previously, we introduced a data-integrated modeling approach we call Methylation-to-Expression Feature Model (M2EFM) (12) and applied it in the case of clear cell renal cell carcinoma (ccRCC). The basis of this approach is to pre-filter loci to those that are differentially methylated between matched tumor and adjacent, pathologically normal tissue and to associate the methylation status of those loci with significant differences in gene expression in the tumor samples. The differentially methylated loci are then called m2eQTL (for methylation-to-expression QTL) and the genes are called m2eGenes. An initial model is built from m2eQTL and m2eGenes with the largest effects to generate risk scores for all samples using Cox-Ridge regression (using the exponential function of the weighted sum of the model features). A final model is built from the risk scores in conjunction with clinical variables and a final risk score is generated from the weighted sum of the features of this model. Here we build M2EFM models of overall survival (OS), distant recurrence-free survival (DRFS), and pathologic complete response (pCR) in breast cancer, where OS is defined as time from initial diagnosis to death from any cause, DRFS is defined as time from diagnosis to distant recurrence, death from cancer, or death from unknown cause, and pCR is defined as a patient being recorded as having a pathologic complete response or a residual cancer burden class of RCB 0/I as in (13).

### Data

A full description of data pre-processing appears in the Supplementary Methods. For this work, we supplied a list of probes to M2EFM from a study of CpG loci involved in epigenetic field effects in breast cancer based on data available in GEO (GSE69914). The loci were delineated CpG sites exhibiting subtle differences in methylation in samples from breast tissue from non-diseased healthy subjects compared to samples from tumor-adjacent normal breast cancer cases, that are also shown to exhibit differential methylation in tumor tissue itself (14). Therefore, they represent regions that may be involved in early loss of regulation in breast cancer.

Aside from the probe list, five different breast cancer cohorts were used in this work. The first dataset is referred to as **TCGA**, was used to train and test an OS model for breast cancer, and consisted of gene expression and DNA methylation profiles created by TCGA, using the Illumina HiSeq 2000 and Illumina Infinium HumanMethylation 450 platforms, respectively. Methylation data were functionally normalized (15) using the *minfi* package (16) in R.

Expression data were TDM normalized, which makes the distributions between microarray and RNA-seq datasets similar, making them more comparable to the validation datasets (17). The samples are described in Table S1. After pre-processing, we retained 1028 samples.

The second dataset is referred to as **Terunuma**, is used for external validation of the OS model, and contains 61 tumor samples with clinical and survival data, downloaded from ArrayExpress (E-GEOD-39004) (18). These data were assayed on the Affymetrix GeneChip Human Gene 1.0 ST Array. They were normalized and then batch corrected in conjunction with the following dataset. A description of the samples is in Table S2.

The third dataset is referred to as **Kao**, is used as a second external validation dataset for the OS model, and contains 327 tumor samples, downloaded from ArrayExpress (E-GEOD-20685) (19). These data were assayed on the Affymetrix GeneChip Human Genome U133 Plus 2.0. A description of the samples appears in Table S2.

The fourth dataset is referred to as **Hatzis1**, was used to train and test a model of DRFS and another model of pCR, and includes HER2-negative breast cancer cases with neo-adjuvant treatment by taxane-anthracycline (followed by endocrine therapy for ER-positive cases) (13). These data were assayed on the Affymetrix Human Genome U133A Array. They were downloaded from GEO (GSE25055), and were normalized and then batch corrected in conjunction with the following dataset. A description of the samples appears in Table S3. After pre-processing we retained 306 samples.

The final dataset is referred to as **Hatzis2**, was used for external validation of both the DRFS and pCR models of (HER2-negative) breast cancer, and includes HER2-negative breast cancer cases with neo-adjuvant treatment by taxane-anthracycline (followed by endocrine therapy for ER-positive cases) (13). These data were assayed on the Affymetrix Human Genome U133A Array. They were downloaded from GEO (GSE25065). A description of the samples appears in Table S3. After pre-processing we retained 182 samples.

In order to reduce categories with very few samples and the number of variables in the model, all AJCC tumor stage classifications were reduced to simple Stage I, II, III, or IV.

### Experimental Design

Three different types of models were built: 1) a model of overall survival (OS), 2) a model of distant recurrence-free survival (DRFS), 3) a model of pathologic complete response (pCR) to neoadjuvant taxane/anthracycline treatment. For all model types, we built 100 different models using different random splits of the data, divided into 70% training and 30% testing data sets, to assess how consistently a good model could be found given different training data. We assessed the performance of the models built with and without clinical variables, with clinical variables alone, and separately to include external validation data. The external validation data could only be tested on models without DNA methylation values, because methylation data were not available (an overview is shown in Fig. 1). For external validation, a final model using all training data was also built.

**Figure 1:**
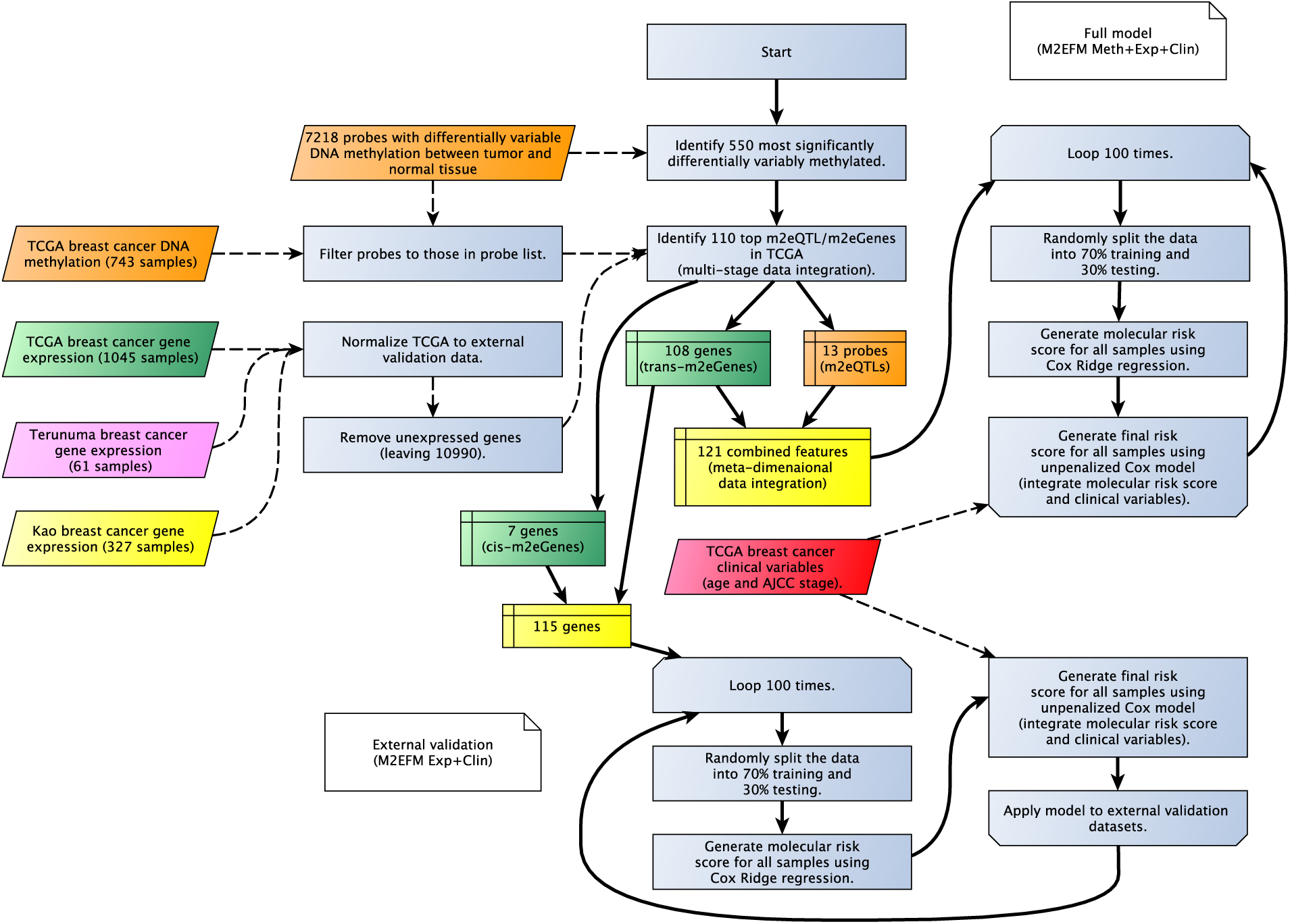
Overview of the M2EFM OS model construction.

M2EFM OS and DRFS models were compared with Cox-Ridge regression, using the *glmnet* package (20) for R, without pre-selection of genes or probes (although probes were pre-filtered by median absolute deviation to reduce them to a number of the most variable probes roughly equal to the number of genes). The M2EFM pCR model was compared to a logistic-Ridge regression model, again using *glmnet*. Furthermore, the M2EFM OS and DRFS models were compared to a model incorporating the PAM50 risk of recurrence score (rorS) (5) with clinical variables. Finally, the M2EFM model was compared to a model built from a gene signature obtained by the network clinical association (NCA) algorithm (21), which is also based on data integration.

Performance of the OS and DRFS models was evaluated using concordance or C-index, a measure of how likely it is, in any given pair of individuals, that the individual with the higher risk score has the event first. For the pCR model, the performance was measured using area under the receiver operating characteristic curve (AUC). Significance tests for comparisons used the two-tailed Wilcoxon signed-rank test.

Clinical variables were chosen using the least absolute shrinkage and selection operator (LASSO) for each outcome. Candidate variables included AJCC stage, patient age at diagnosis, ER status, PR status, HER2 status, and PAM50 subtype (calculated using the *genefu* package for R) (22). Selected variables include AJCC stage and patient age at diagnosis for all outcomes, although stage was reduced to simply presence or absence of stage III cancer for DRFS and pCR outcomes due to the absence of stage IV cases and small number of stage I cases for these outcomes.

The m2eGenes were used to analyze the functional relevance of our approach using the online tool WEB-based GEne SeT AnaLysis Toolkit (WebGestalt 2017) (23). Using WebGestalt, we performed protein network topology-based analysis and enrichment analysis for GO biological process terms. Furthermore, we tested the enrichment of m2eGenes in a database of genes with causal cancer connections, the Catalog of Somatic Mutations in Cancer (COSMIC) Cancer Gene Census (24), using Fisher’s exact test. Finally, we tested the enrichment of genes in our signature in genes known to be targeted by breast cancer drugs listed in the DrugBank database (25), again using Fisher’s exact test.

## Results

We applied M2EFM to the TCGA dataset to select dysregulated probes and genes for our models. The lists of probes and genes selected by M2EFM are shown in Table S4.

### Overall Survival – Meta-dimensional Models Without External Validation Data

Results of the meta-dimensional double-integrated model, (Fig. 1 and fully described in the Supplemental Methods) are shown in Fig. 2A and Table S5, allowing a comparison to how frequently equivalent strength results could be obtained from random gene sets. Across the 100 random splits of the training and testing data, the double integrated model (M2EFM Meth+Exp+Clin) had the highest median C-index at .790, followed by PAM50 rorS values for the samples and clinical variables (rorS+Clin) at .775, then the clinical variables only model (Clin) at .753, and finally the Cox-Ridge regression with an additional regression integrating unpenalized clinical variables (Cox Meth+Exp+Clin) at .728. The C-index scores across the splits were statistically significantly greater for the M2EFM Meth+Exp+Clin model than rorS+Clin, Clin, or Cox Meth+Exp+Clin (p = 1.84e-08, 1.26e-12, and 3.23e-16 respectively). For models built from only the molecular risk score, M2EFM Meth+Exp had the highest median C-index although scores tended to be lower than models incorporating clinical data. M2EFM Meth+Exp results were significantly greater than rorS and Cox Meth+Exp (p = 3.44e-11, and 1.13e-02 respectively).

**Figure 2:**
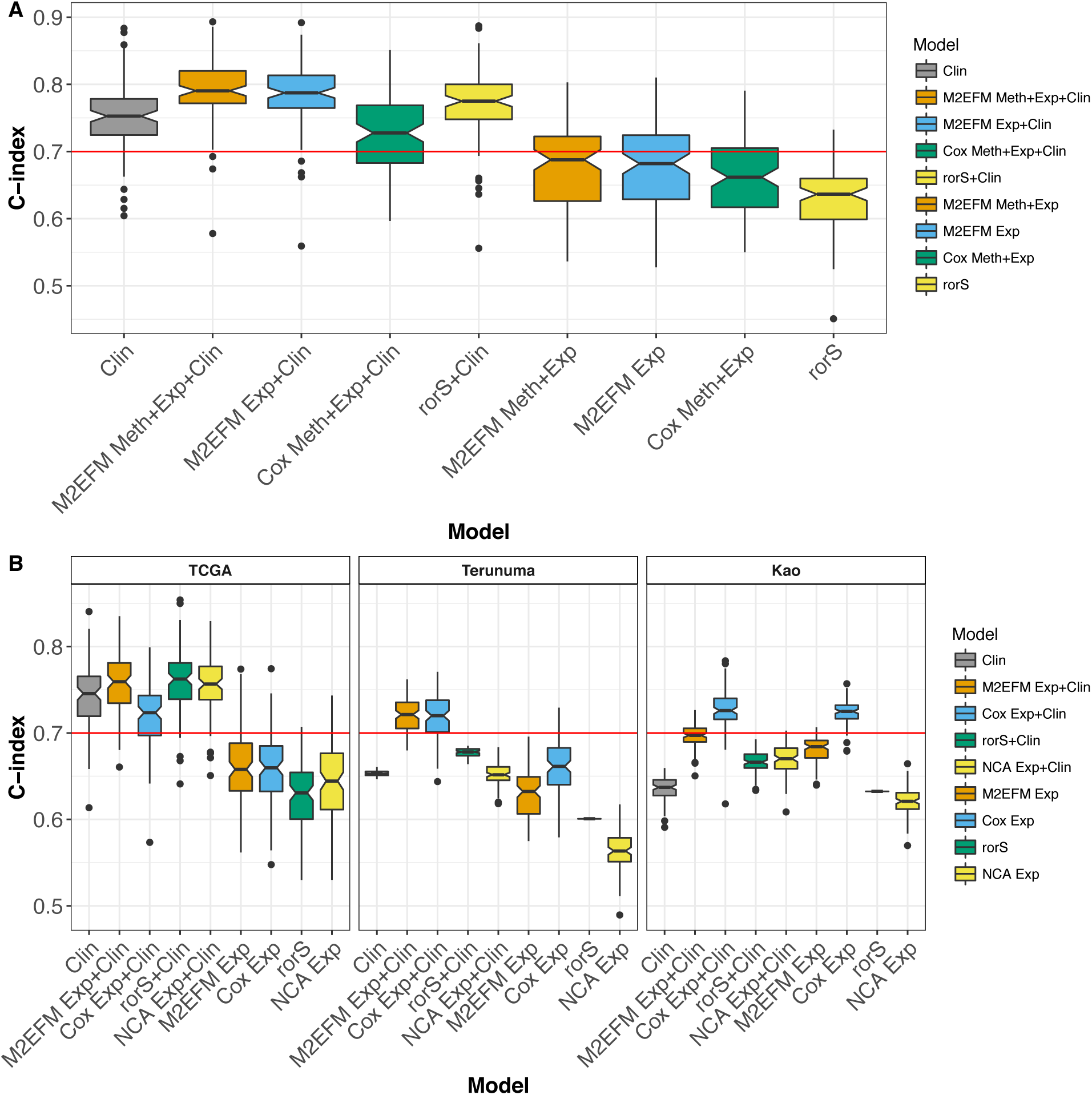
C-index for full overall survival models across 100 random splits of TCGA data into training and testing data for the various approaches. M2EFM models are the most consistent performers, achieving a median C-index of at least .7 in both internal and external validation data in all cases. A) The M2EFM model with gene expression, DNA methylation, and clinical data (M2EFM Meth+Exp+Clin) was statistically significantly stronger than any other model. B) The models with external validation data show that only the M2EFM and Cox-Ridge models consistently validate at a high median C-index.

### Overall Survival - Expression Only Models with External Validation Data

To allow external validation of our models using other datasets we next focused on breast cancer OS, building models using only expression data, with the m2eQTL discovered previously. In such cases, we substitute the *cis*-m2eGenes (genes proximal to the differentially methylated loci) for the associated methylation values. We also added a model built from a gene signature discovered using NCA (except for two genes: *HES5* and *NKIRAS1*, which were not in our profiles).

The median C-index for each model on each dataset is shown in Table S6, and the distribution of C-indices is shown in Fig. 2B. On TCGA, the median C-index for rorS+Clin was the highest (.762), followed closely by M2EFM Exp+Clin (.759) and NCA Exp+Clin (.757), with the other models further behind. The differences were not significant for rorS+Clin compared to M2EFM Exp+Clin (p = 6.76e-02) but were for NCA Exp+Clin, Clin, and Cox Exp+Clin (p = 3.99e-02, 3.26e-11, and 5.80e-13 respectively). However, the M2EFM model was the most consistent performer over all the validation data. On the Terunuma external validation data, M2EFM Exp+Clin validated better than any other model, with a median C-index of .721, followed by Cox Exp+Clin at .720, while the other models trailed behind. The M2EFM Exp+Clin C-index scores were not significantly greater than Cox Exp+Clin (p = 8.17e-01), but were significantly greater than rorS+Clin (p < 2.20e-16) and NCA Exp+Clin (p < 2.20e-16). In the Kao data, the best performer was Cox Exp (.726), followed by Cox Exp+Clin (.725), then M2EFM Exp+Clin (.697), followed by the other models. The C-index scores for Cox Exp were significantly stronger than M2EFM Exp+Clin (p = 3.14e-16), but M2EFM Exp+Clin scores were still significantly stronger than either rorS+Clin (p < 2.20e-16) or NCA Exp+Clin (p < 2.20e-16).

### Full Training Data Model for Overall Survival

We used the full training data set (TCGA), to build a final overall survival model and validated it on Terunuma and Kao. For Terunuma, the C-index was .724, 95% CI [0.499, 0.874]. For Kao, the C-index was 0.703, 95% CI [0.582, 0.800]. These scores were very similar to their median C-index scores. The hazard ratios for this model are given in Table S7. We further divided the samples into three groups based on their risk score, with low risk below the 25^th^ percentile, medium risk from the 25^th^ to the 75^th^ percentile, and high risk above the 75^th^ percentile, and created Kaplan-Meier plots of the survival of the risk groups in each dataset (Fig. S1). Hazard ratios comparing risk groups for the risk score are shown in Table 1.

**Table 1:**
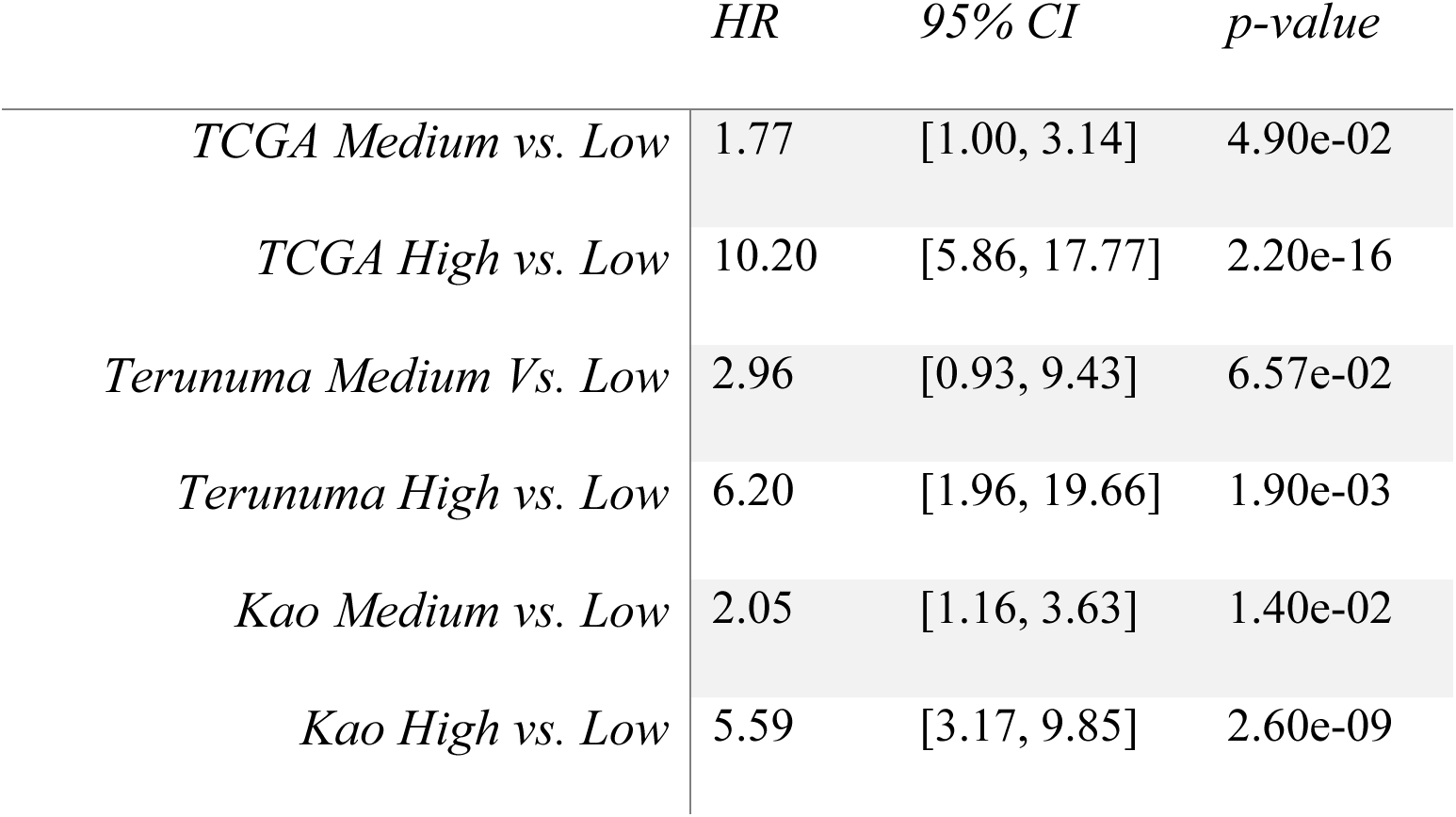
*Hazard Ratios for Overall Survival Model Risk Groups*

We generated calibration curves for the proportional hazards models based on the full training data model (Fig. S2), using the *pec* package for R (26), as well as the integrated Brier score (IBS) (27), a quantitative measure of calibration or prediction error. For M2EFM Exp+Clin, the IBS on Kao was .060 and on Terunuma was .166. For Cox Exp+Clin, the IBS on Kao was .055 and on Terunuma was .161, indicating that the Cox Exp+Clin models were better calibrated over all time points.

### ER Status

Most prognostic markers for breast cancer have limited utility for patients with ER-negative tumors (28). Unfortunately, ER status was not annotated in the Kao data, and there were too few samples in Terunuma to effectively test the prognostic utility of our marker on ER-negative patients. Therefore, we inferred the ER, PR, and HER2 status of samples in Kao by clustering samples using expression probes associated with each receptor status using a Gaussian mixture model to capture each bimodal distribution (28, 36). We then assessed the relative prognostic power of the M2EFM models in samples that varied by ER or triple negative status. The overall C-index was .703, 95% CI [0.582, 0.800], but for ER-positive patients it was .665, 95% CI [.489, .806], for ER-negative patients it was .727, 95% CI [.558, .849], and for triple negative patients it was .682, 95% CI [.388, .879].

### Subtypes

Given that the covariates in our model are unpenalized, we needed to select a limited number of clinical covariates to avoid overfitting the model. This resulted in models without PAM50 intrinsic subtypes, ER, PR, or HER2 status, which are all known to be associated with breast cancer prognosis. Therefore, we regressed the M2EFM molecular risk score (calculated for all TCGA samples) on these variables and calculated the variance inflation factor (VIF), a commonly used measure of the multicollinearity between variables. The VIF was approximately 1.20. A VIF between 1 and 5 is generally regarded to imply only moderate multicollinearity.

Therefore, the molecular risk score from M2EFM contains prognostic information independent of breast cancer subtype.

### Distant Recurrence-Free Survival (DRFS) Models

On both the testing data and the external validation data, M2EFM was the most prognostic model of DRFS (Fig. 3 and Table S8). On the testing data, the M2EFM Exp+Clin had a median Cindex score of .722, followed by rorS+Clin (.707), then M2EFM Exp (.705), and Cox Exp+Clin (.701), with the other models below .7. In the external validation data, M2EFM Exp+Clin and M2EFM Exp were the most prognostic at .752 and .741 respectively, although Cox Exp+Clin and Cox Exp were strong at .735 and .729 respectively.

**Figure 3:**
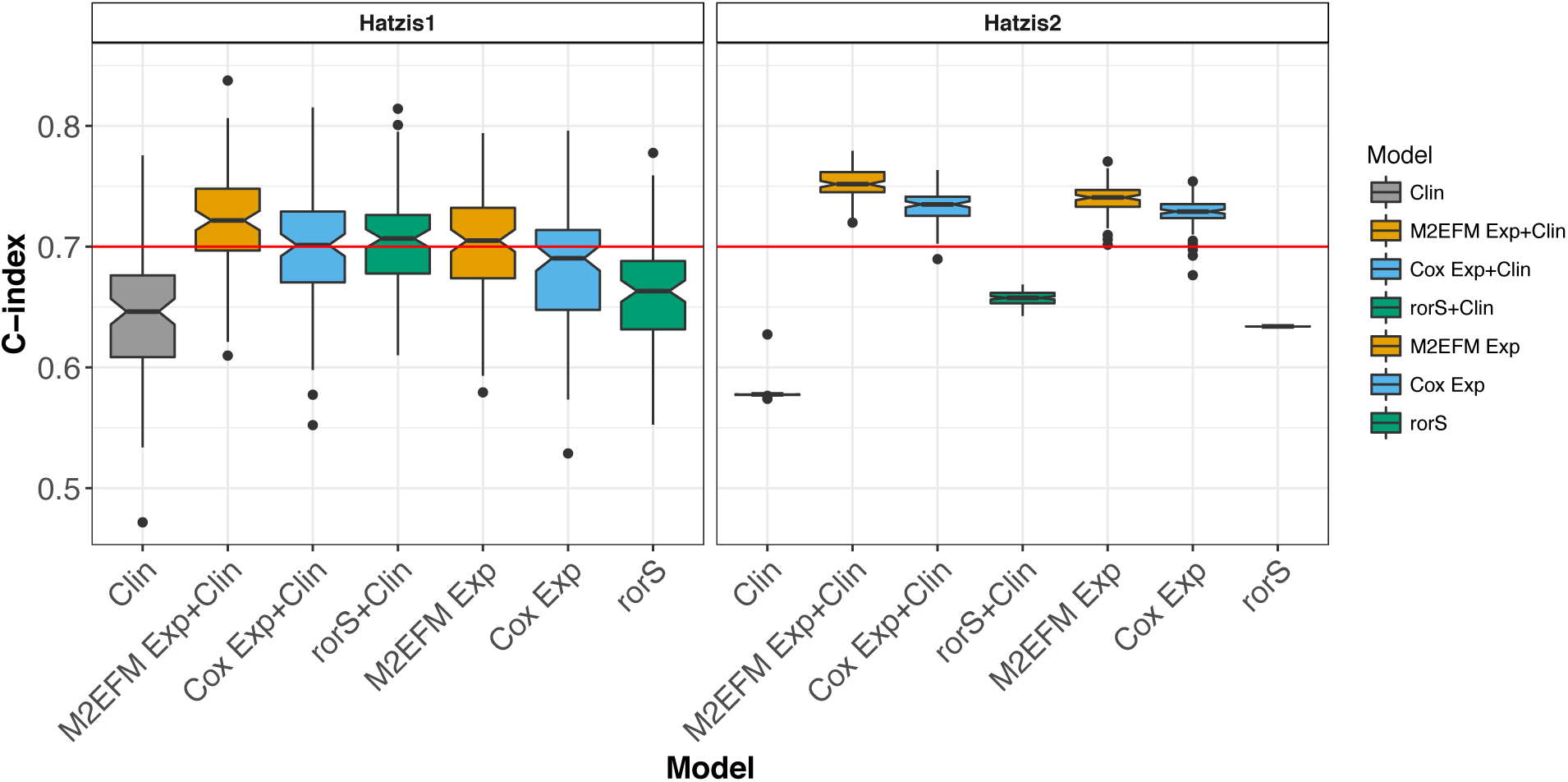
C-index for expression-only DRFS models across 100 random splits of Hatzis1 data into training and testing data for the various approaches, which were then validated on Hatzis2. The M2EFM models were the best performing on both testing and external validation data.

Finally, an M2EFM Exp+Clin model trained on the full training data (Hatzis1) achieved a Cindex of .766, 95% CI [.604, .875] on the external validation data (Hatzis2). The hazard ratio for this model are given in Table S9. Calibration curves at 3 years are shown in Fig. S3. The IBS score for M2EFM Exp+Clin on Hatzis2 was 0.079. For Cox Exp+Clin it was 0.083, demonstrating that the M2EFM Exp+Clin model is better calibrated.

#### Negative Predictive Value for Node-Negative Cases

One interest in genomic cancer biomarkers is for determining cases with a low risk of recurrence who may not benefit from adjuvant chemotherapy (37, 38). For these cases, it is important that the marker has a high negative predictive value (NPV). Therefore, we assessed the NPV of the M2EFM model in node-negative samples using the *timeROC* package for R, which uses a time-dependent AUC, which can handle censored survival times. A cutoff for the risk score was determined in Hatzis1 that equalized the sensitivity and specificity in predicting DRFS for node-negative cases at 3 years (the median follow-up in these data). This cutoff was applied to node-negative cases in Hatzis2 to determine the positive predictive value (PPV) and NPV. At the 3-year follow up point, the PPV was .49 and the NPV was .95. We predicted 39 node-negative cases would not have a distant metastasis by 3 years and we were correct for 36. We predicted 12 node-negative cases would have a distant metastasis and were correct for 7.

### Pathologic Complete Response (pCR) Models

The final set of models are for pCR following neoadjuvant treatment with Taxane/Anthracycline in HER2-negative breast cancer. Results are shown in Fig. 4 and Table S10. For these data, the clinical models were not very informative. The reference logistic-Ridge regression model performed best on the testing data, with Logistic Exp+Clin scoring a median AUC of .767 and Logistic Exp at .763 compared to M2EFM Exp+Clin with a median AUC of .739 and M2EFM Exp at .733. Nevertheless, M2EFM performed best on the external validation data, with M2EFM Exp+Clin at a median AUC of .716 and M2EFM Exp at .727 compared to Logistic Exp+Clin at a median AUC of .707 and Logistic Exp at .709.

**Figure 4:**
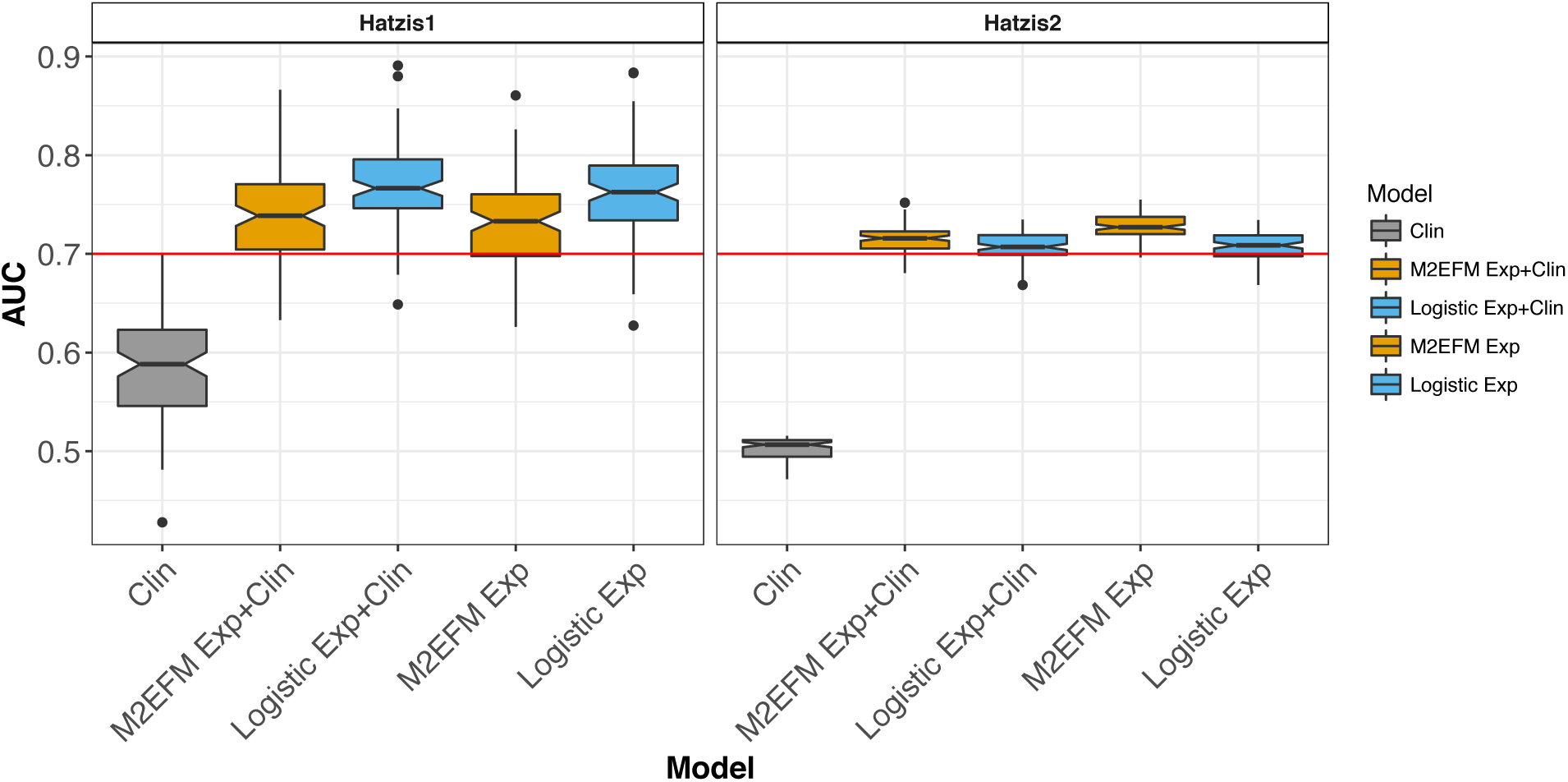
AUC for expression-only pCR models across 100 random splits of Hatzis1 data into training and testing data for the various approaches, which were then validated on Hatzis2. Although Logistic-Ridge models performed better on the internal validation models, M2EFM models performed better on external validation data.

Finally, an M2EFM Exp+Clin model trained on the full training data (Hatzis1) achieved an AUC of .720, 95% CI [.639, .802] on the external validation data (Hatzis2). The odds ratios for this model are given in Table S11. Calibration curves appear in Fig. S4. The mean Brier score for M2EFM Exp+Clin on Hatzis2 was .184 and for Cox Exp+Clin it was .202, showing that M2EFM Exp+Clin is better calibrated.

### Functional Analysis

#### Protein Network Topology-Based Analysis

Of the 115 m2eGenes, 55 were found to interact in a sub-network of protein-protein interactions using WebGestalt. The sub-network included 65 genes in all and was enriched for 184 biological process terms, demonstrating that many of the genes in our signature are functionally related. The top result was for lymphocyte activation (p = 2.50e-12), which included 23 genes from the sub-network and is about 8.96 times more overlap than would be expected by chance. The subnetwork is shown in Fig. S5, visualized using Cytoscape (32).

#### Biological Process, Disease, and Drug Enrichment

Using Gene Ontology enrichment testing, our gene set shows a clear enrichment for immune functions (Fig. 5). The results further demonstrate that expression tends to be repressed in these pathways in the samples with the highest M2EFM risk scores and more moderately repressed in samples with medium M2EFM risk scores.

**Figure 5:**
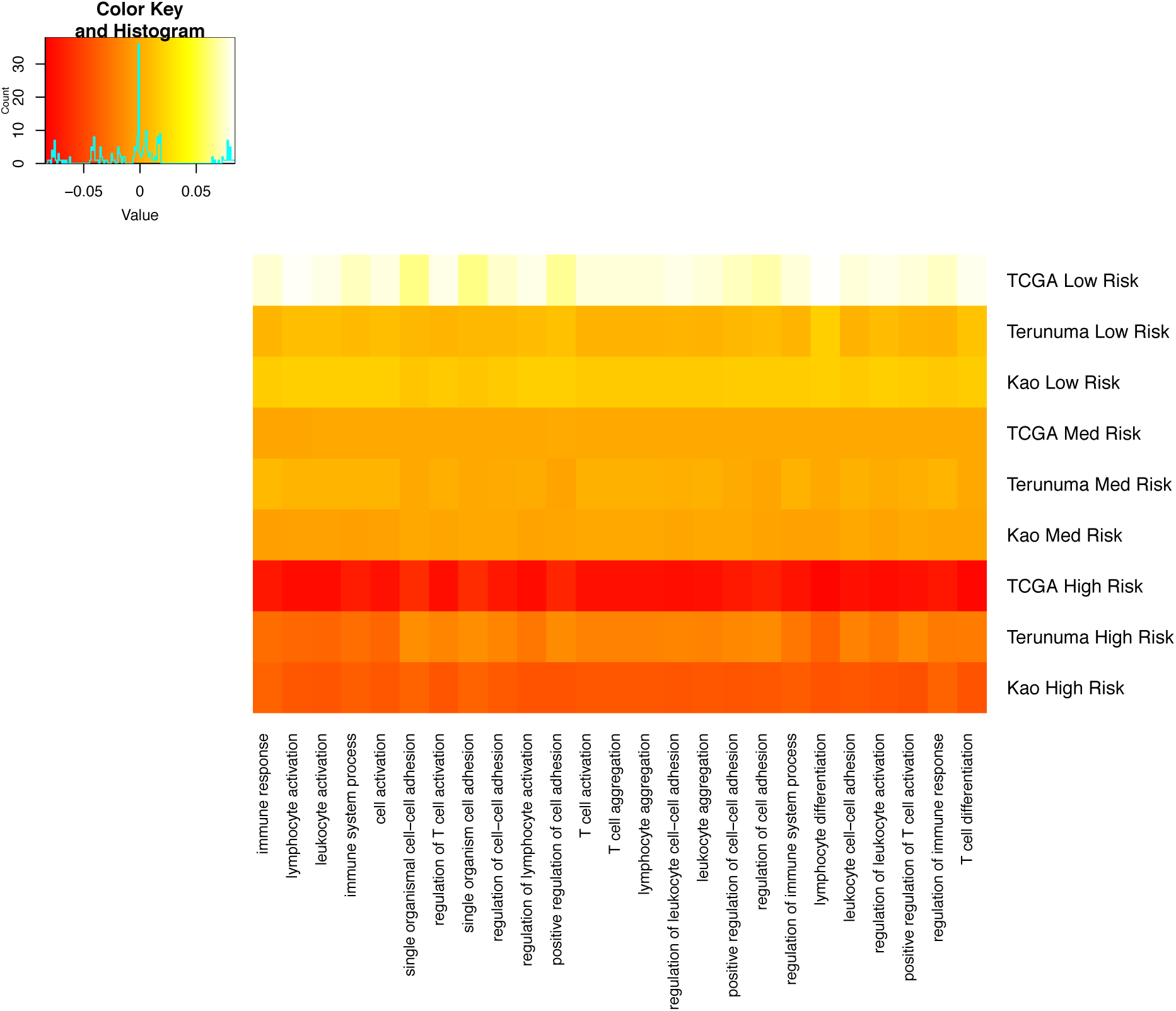
Top 25 biological process pathways enriched for genes in the M2EFM gene signature. The pathways are composed primarily of immune-related functions. The heat map shows the mean relative expression of genes from our signature in each pathway for each overall survival dataset. The data are subset into samples predicted by M2EFM to be at low, medium, or high risk. There is a trend toward repression of the pathways as the risk score goes from low to high.

Many of the genes in our gene signature are already known to be associated with breast cancer (such as *ESR1* (33) and *EGFR* (34)). We calculated the enrichment of genes in our signature in the Catalog of Somatic Mutations in Cancer (COSMIC) Cancer Gene Census (24), downloaded on 9/13/2016. Enrichment would suggest a causal link between genes in our signature and the progression of the disease. There were 601 genes in the catalog, 515 of which were in our gene expression profiles. The enrichment for causally implicated cancer genes in our gene signature was about 4 times greater than expected by chance (OR: 3.87, p = 6.09e-06). The following 18 genes used in our M2EFM models appear in the Cancer Gene Census: *AFF3*, *BCL11A*, *CD79A*, *CD79B*, *EGFR*, *ERBB4*, *ESR1*, *FOXA1*, *GATA3*, *IRF4*, *ITK*, *LCK*, *MYB*, *POU2AF1*, *PRF1*, *PTPRC*, *RET*, *TNFRSF17*.

We also examined the known targets of FDA approved breast cancer drugs using the DrugBank database version 5.0 (25). There were 43 targets among the genes in our expression profiles. Of these, 4 genes were found in our signature (*EGFR*, *ESR1*, *MAPT*, and *PRKCB*). Thus, our signature was enriched for known targets of breast cancer treatments 10 times more than would be expected by chance (OR: 10.00, p=1.03e-03). These drugs included paclitaxel (*MAPT*), raloxifene (*ESR1*), toremifene (*ESR1*), fulvestrant (*ESR1*), trastuzumab (*EGFR*), tamoxifen (*ESR1*, *PRKCB*), docetaxel (*MAPT*), and lapatinib (*EGFR*).

## Discussion

In this work, we have tried to address common shortcomings in molecular biomarker studies, have applied the REMARK guidelines in our assessment (35), and validated a data-integrated approach to creating prognostic and predictive cancer models. The M2EFM approach shows remarkable promise for developing prognostic and predictive cancer biomarkers using both clinical and molecular data. For overall survival, distant recurrence free survival, and pathologic complete response in breast cancer it achieved a median C-index or AUC of at least .7 in all internal and external validation datasets (including 5 different cohorts). The fact that our model can reliably attain this level of discrimination across platforms and populations is promising indeed.

Unsurprisingly, one of our comparison methods, a modified Cox-Ridge approach, performed about as well as M2EFM in terms of discriminative power. Given that M2EFM depends on Cox-Ridge itself, those models contain a superset of the predictors in M2EFM. Nevertheless, M2EFM models in many cases outperform the Cox-Ridge models, and more importantly, M2EFM models depend on only 115 genes, as opposed to 10990 in the Cox-Ridge models. Therefore, clinical use for Cox-Ridge would require a transcriptome-wide gene expression assay, and the situation would be even more extreme if the model with methylation values was used. Additionally, for DRFS and pCR, M2EFM was better calibrated than Cox-Ridge, suggesting that Cox-Ridge would not be as useful clinically. We also demonstrated that M2EFM has a high NPV for node-negative cases, while maintaining a meaningful PPV, suggesting M2EFM might be useful for identifying patients who could avoid adjuvant chemotherapy. Finally, M2EFM has functional relevance, increasing the possibility that it could serve as the basis for clinical tests that determine eligibility for treatment.

Thus, using three different outcomes and five different data sets, we maintained a consistent level of accuracy. In conjunction with our prior study of M2EFM applied to ccRCC (12), these results consistently show that integrating both molecular and clinical data results in more prognostic models than either data type alone. Our results strongly suggest that a data-integrated approach that models genes affected by regulatory changes in cancer is more reliable than many extant approaches. Furthermore, our results suggest our method builds a model that is prognostic in the case of ER-negative samples. This is in contrast to most breast cancer prognostic biomarkers, including the PAM50 rorS.

Our results also show that our approach does more than build a reliable prognostic model: it identifies biologically relevant pathways and, importantly, possible therapeutic targets and genes involved in cancer progression. Our gene signature contains genes that are functionally involved in cancer and that are targeted by current breast cancer treatments. Given recent advances in immunotherapy, it is interesting to note the repression of immune related genes in our signature for patients predicted to be at high risk.

Although we provide more evidence of the efficacy of M2EFM than most computationally derived molecular biomarker studies, there remains one limitation before attempting to translate these results to the clinic: calibration of the model to a single platform. To obtain external validation data, we performed cross-platform normalization. Having established the utility of the M2EFM approach, it remains to train the model on a high-quality dataset generated on the platform that will be used for any clinical assays.

M2EFM can predict risk based on data integrated from multiple sources, taking advantage of tried-and-true prognostic factors while incorporating data on relevant relationships between molecular data types at the individual level. For breast cancer, this resulted in models based on several genes with known functional associations to cancer progression and which are targeted by currently available drugs, and therefore, may result in a useful tool for clinical decision support. This also suggests that other genes in the model should be examined as possible targets, because many of the genes are functionally connected. Given the strength of our findings, we believe that further development of this technique and application to other cancers is warranted.

## Availability

M2EFM is available as a package for R at https://github.com/jeffreyat/M2EFM.

